# Characterising cortical perturbation-recovery dynamics using TMS-EEG: The Brain Dynamic Metric Toolbox

**DOI:** 10.64898/2026.10.30.763068

**Authors:** Wei Qin, Andre Peterson, Anthony Burkitt, Paul B Fitzgerald, Philippa J Karoly, Jodie Naim-Feil

## Abstract

**Objective:** To adapt dynamical measures used in computational and experimental perturbation models to human transcranial magnetic stimulation-electroencephalography (TMS-EEG) and determine whether analysis of the extended post-stimulation trajectory reveals additional features of the cortical response to long-interval intracortical inhibition (LICI).

**Methods:** Single-pulse and paired-pulse LICI TMS-EEG responses from the bilateral dorsolateral prefrontal cortex were re-analysed in 16 healthy adults. At the stimulation site, 75 single-pulse and 75 paired-pulse trials were delivered; paired pulses comprised conditioning and test stimuli separated by 100 ms. Responses from 20 frontocentral EEG channels were analysed from 50–950 ms following TMS using the open-source Brain Dynamic Metric Toolbox. Perturbative Integration Latency Index (PILI) indexed cortical resilience via response energy and recovery, Autocorrelation Function Width (ACFW) captured temporal memory, and Lyapunov Exponent (LE) indicated the dynamical stability. Linear mixed-effects models examined temporal, spectral, and time-frequency effects from the measured metrics.

**Results:** LICI altered the temporal evolution of the dynamical measures. PILI distinguished conditions in the earliest post-stimulation window before converging, whereas ACFW differences persisted across a broader early interval. LE showed weaker and less consistent discrimination. Spectral effects were most evident for ACFW, while time-frequency analyses revealed later, frequency-specific LICI effects for PILI and ACFW but not LE.

**Conclusions:** The extended post-stimulation trajectory revealed LICI-related differences in cortical resilience and temporal memory with distinct temporal and spectral profiles, while effects on dynamical stability were less consistent.

**Significance:** The Brain Dynamic Metric Toolbox provides a reproducible framework for quantifying perturbation-recovery dynamics in TMS-EEG and demonstrates that the extended perturbational response contains information about LICI not captured by discrete early response measures.

**Graphical Abstract:** 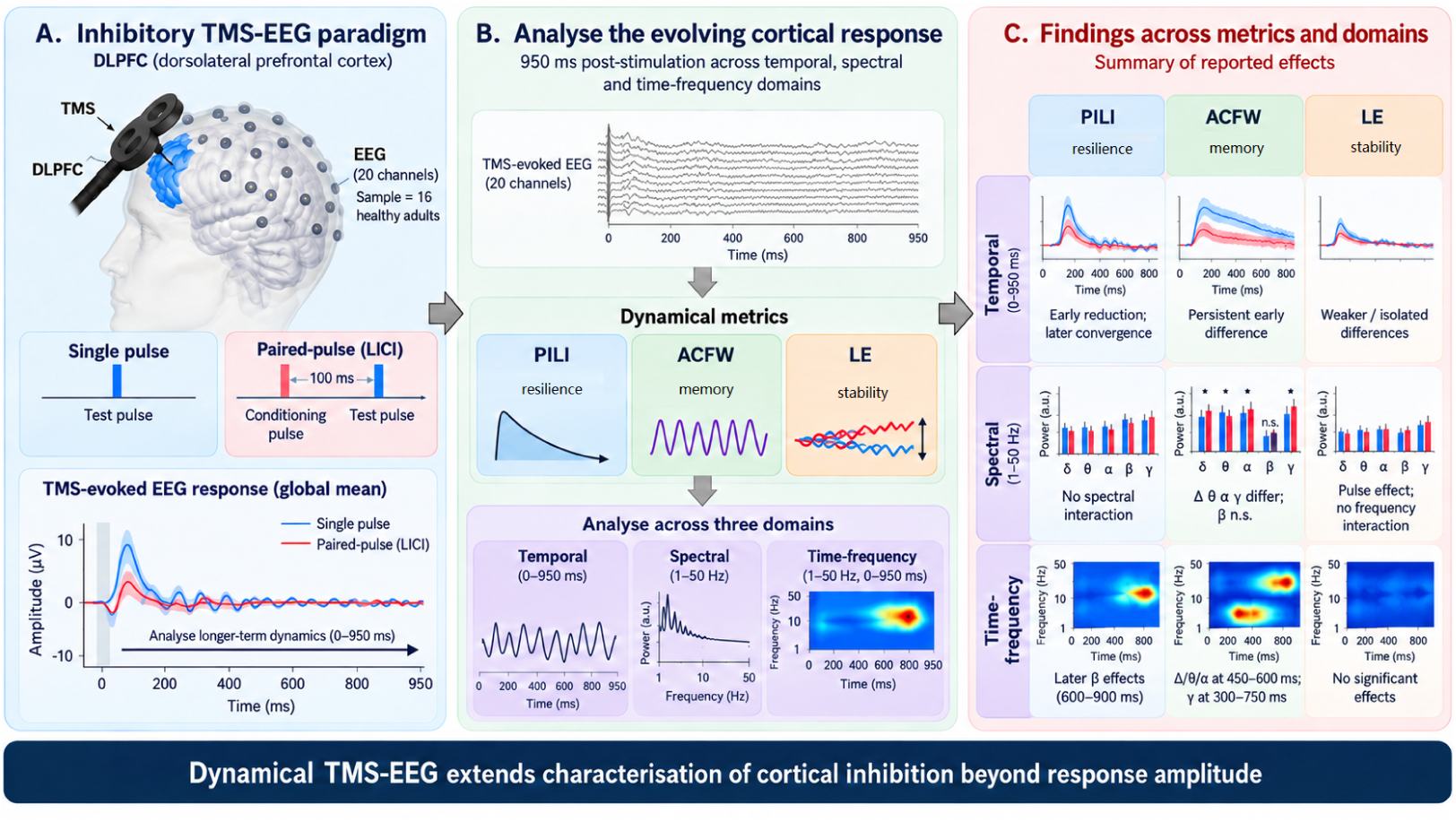

Responses to single-pulse transcranial magnetic stimulation with electroencephalography (TMS–EEG) and paired-pulse stimulation using long-interval intracortical inhibition (LICI) were compared in 16 healthy adults following stimulation of the dorsolateral prefrontal cortex. Three dynamical metrics, the Perturbative Integration Latency Index (PILI), Autocorrelation Function Width (ACFW) and Lyapunov Exponent (LE), characterised the post-stimulation response across temporal, spectral and time-frequency domains over 950 ms. Inhibitory conditioning was associated with early alterations in cortical resilience, more persistent changes in temporal memory and later frequency-specific effects. LE differences were less consistent. Together, these measures provide complementary characterisation of the evolving response to inhibitory conditioning and are implemented in the user-friendly, open-source Brain Dynamic Metric Toolbox to support reproducible analysis and further physiological and clinical validation.

**Highlights:**

- Dynamical metrics quantify a perturbation-recovery framework for inhibitory TMS-EEG
- Analysis beyond 300 ms post TMS revealed additional dynamics of paired-pulse inhibition
- Dynamical metrics captured distinct temporal and spectral inhibitory responses
- Extended dynamics may help characterise inhibitory processes in brain disorders
- Open-source toolbox enables reproducible perturbation-recovery analysis

## 1. Introduction

Transcranial magnetic stimulation combined with electroencephalography (TMS-EEG) provides a non-invasive index of cortical excitability, inhibition, and effective connectivity by measuring the brain’s electrical response to a brief magnetic perturbation (Ilmoniemi et al., 1997; Komssi and Kähkönen, 2006; Rogasch and Fitzgerald, 2013; Ziemann et al., 2026). This perturbation approach allows the cortical response to be characterised in vivo with millisecond temporal resolution, including the propagation of evoked activity following stimulation. TMS-EEG is a valuable tool for probing circuit pathophysiology and treatment-relevant brain dynamics across psychiatric and neurological disorders (Tremblay et al., 2019; Farzan, 2024; Ziemann et al., 2026).

Paired-pulse paradigms, such as long-interval intracortical inhibition (LICI), apply a suprathreshold conditioning pulse to suppress the response to a subsequent test pulse approximately 100 ms later, with this suppression commonly interpreted as reflecting cortical inhibitory mechanisms, including processes consistent with GABA_B_ receptor-mediated inhibition (McDonnell et al., 2006; Daskalakis et al., 2008; Rogasch et al., 2013). When combined with EEG, LICI can be measured directly from cortical evoked activity, enabling assessment of inhibition in non-motor regions such as the dorsolateral prefrontal cortex (DLPFC) (Daskalakis et al., 2008; Fitzgerald et al., 2009; Rogasch et al., 2015). Mapping altered cortical reactivity within these regions is particularly relevant for clinical populations in which frontal inhibition and excitation-inhibition balance are disrupted and are difficult to index non-invasively in humans. TMS-EEG studies have reported altered frontal excitability, inhibitory physiology, and network organisation across psychiatric, neurological, and neurodegenerative disorders (Farzan et al., 2010; Radhu et al., 2013; Naim-Feil et al., 2016; Sun et al., 2016; Benussi et al., 2017; Naim-Feil et al., 2022; Fatih et al., 2021).

Conventional TMS-EEG analyses typically summarise the evoked response using early post-stimulation components. Although informative, these measures may not fully capture how the response evolves over a broader post-stimulation interval (i.e., *>* 300 ms), including response resilience, temporal memory and spectral organisation. Disorders involving altered excitation-inhibition balance may differ not only in the magnitude of the evoked response, but also in the temporal evolution of cortical activity following perturbation. Quantifying these recovery dynamics may therefore provide an additional approach for characterising the effects of paired-pulse inhibition beyond suppression at selected latencies or electrodes. This rationale is supported by converging evidence from intracranial recordings and computational modelling, which indicates that TMS-evoked activity is not confined to isolated local responses but evolves through distributed networks over several hundred milliseconds (Pigorini et al., 2011; Mutanen et al., 2013; Momi et al., 2021, 2023, 2025; Tsang et al., 2026). Characterising the response across an extended temporal window may therefore provide additional information about its resilience and temporal memory.

To address this gap, dynamical systems theory offers a complementary framework for quantifying perturbation-recovery dynamics. Rather than reducing the response to discrete peaks, these metrics treat the brain as a nonlinear dynamical system and characterise different properties of the evolving cortical response. Measures following a perturbation can characterise the resilience and recovery of activity over time; autocorrelation captures temporal memory; and Lyapunov-based measures quantify trajectory divergence as an index of dynamical stability. These features are potentially informative because inhibitory conditioning may be expressed not only through changes in response amplitude, but also through changes in cortical resilience and temporal memory. These metrics may therefore be particularly well-suited to paired-pulse paradigms, where inhibition may alter both the magnitude of the evoked response and its evolving temporal structure, providing a broader characterisation of LICI-related response dynamics than amplitude alone. Related perturbational and dynamical frameworks have informed studies of consciousness, cortical complexity, and stimulation-induced recovery (Massimini et al., 2005; Casali et al., 2013; Deco et al., 2018; Jobst et al., 2017; Momi et al., 2023), providing a basis for extending these approaches to paired-pulse inhibition.

To standardise quantification of perturbation-recovery dynamics, we developed the open-source Brain Dynamic Metric Toolbox to implement three complementary metrics for the analysis of TMS-EEG data: the Perturbative Integration Latency Index (PILI) (Jobst et al., 2017) adapted to characterise the resilience of the post-stimulation response; the Autocorrelation Function Width (ACFW) (Scheffer et al., 2009), indexing temporal memory; and the Lyapunov Exponent (LE) (Iasemidis et al., 1990), indexing dynamical stability. The present study applied these metrics to single-pulse and paired-pulse LICI data acquired from the DLPFC in healthy adults (Naim-Feil et al., 2016), extending analysis to 950 ms post-stimulation across temporal, spectral, and time-frequency domains. We hypothesised that paired-pulse stimulation would alter cortical resilience and reduce the temporal memory of the post-stimulation responses relative to single-pulse stimulation, consistent with inhibitory suppression of cortical dynamics. Potential changes in dynamical stability were examined to determine whether this aspect of the response differed following inhibitory conditioning. The open-source implementation supports replication and provides a reproducible framework for subsequent physiological and clinical validation to facilitate future biomarker development in clinical populations.

## 2. Methods

This data was collected as a collaborative effort between Monash Alfred Psychiatry Research Centre and Turning Point Alcohol and Drug Centre. Ethics approval was granted by the Alfred Human Subjects Research and Ethics Committee. Active recruitment and participation occurred from October 2010 through September 2011. All participants signed an informed consent form and were notified that participation was voluntary and they were able to withdraw at any time without prejudice. This study is a re-analysis of data previously published (Naim-Feil et al., 2016, 2022). The present study expands on these earlier studies by 1. Expanding the timeframe of analysis to 950 ms, 2. Applying novel dynamical metric analysis, and 3. Using this data to build a user-friendly TMS-EEG analysis toolbox.

### 2.1. Sample characteristics

Sixteen healthy subjects (mean age = 32 years, SD = 6 years; 8 males and 8 females) were recruited for the study through posters and local advertisements. Exclusion criteria included any documented psychiatric illness, previous or current history of alcohol/drug use disorder, head injury, or history of seizures. At screening, participants completed a demographics questionnaire, assessment of cognitive functioning (Wechsler, 2001), depressive symptoms (Steer et al., 1987) and alcohol dependence scales (Anton et al., 1996; Stockwell et al., 1979), with all results for the cohort within the normal range. Participants received 40 AUD reimbursement for participation.

### 2.2. Transcranial Magnetic Stimulation

A figure-of-eight cooled coil was connected to the MagPro R30 stimulator with a MagOption unit (Magventure, Denmark) with Biphasic TMS pulses applied to the cortex. The coil was angled approximately 45*^◦^* from the midsagittal line and held perpendicular to the presumed direction of the central sulcus. To identify the optimal stimulation parameters, active TMS was first applied to the left motor cortex to index the Resting Motor Threshold (RMT). RMT was set as the minimum stimulus intensity needed to elicit peak-to-peak motor-evoked potentials *>* 50 µV in *>* 5/10 trials (Rossini et al., 1994). RMT was measured using disposable disc electrodes placed across the contralateral first dorsal interosseous (FDI) muscle, with electromyography (EMG) response recorded via Signal software (Cambridge Electronics Design, CED Micro 1401 mk II analogue-to-digital converting unit, Cambridge, UK). RMT was determined for each participant (Mean = 52, SD = 8). Active Motor Threshold (AMT) indexes the minimum stimulation intensity (during FDI muscle contraction i.e., continuously pressing index finger against a spring), required to induce peak-to-peak MEPs of *>* 100 µV in 3/5 consecutive trials. For the 1-mV peak-to-peak measure, the stimulator intensity is adjusted until peak-to-peak MEP of approximately 1mV over 10 consecutive trials is identified. The 1-mV peak-to-peak measure was applied over the EEG cap and measured via the contralateral FDI muscle contraction and these parameters were used to determine the appropriate stimulation intensity for application of frontal TMS-EEG.

Paired-pulse TMS was then applied to both the right and left dorsolateral prefrontal cortex (DLPFC) with the order of stimulation site (left or right) counterbalanced across participants. For frontal stimulation, TMS-cortical activity was measured via a custom-made 24-channel EEG cap with signals recorded using the Synamps2 EEG system (Compumedics Neuroscan, TX, USA) and electrodes (sintered Ag/AgCl) positioned according to standard 10–20 design (EASYCAP GmbH, Germany) and referenced to an electrode posterior to the Cz electrode. EEG signals (DC) were acquired at sampling rate of 20 kHz and filtered (low pass filter 3500 Hz). Following the TMS pulse, recharging capacitors had 1-s delay to minimize TMS-artefacts. All participants had headphones with white noise (95 dB) throughout the session to reduce the presence of auditory artefacts.

### 2.3. Measurement of Long interval intracortical inhibition (LICI)

Long interval intracortical inhibition (LICI) is a paired-pulse inhibitory paradigm. For this, two suprathreshold pulses are applied to the frontal cortex based on the stimulation intensity determined from the 1mV peak-to-peak measure (from motor stimulations). The suprathreshold conditioning pulse is applied, after an inter-stimulus interval (ISI) of 100 ms, this is then followed by a suprathreshold test pulse. The conditioning pulse suppresses this cortical response to the test stimulus. This process is described in Figure 1.

**Figure 1:**
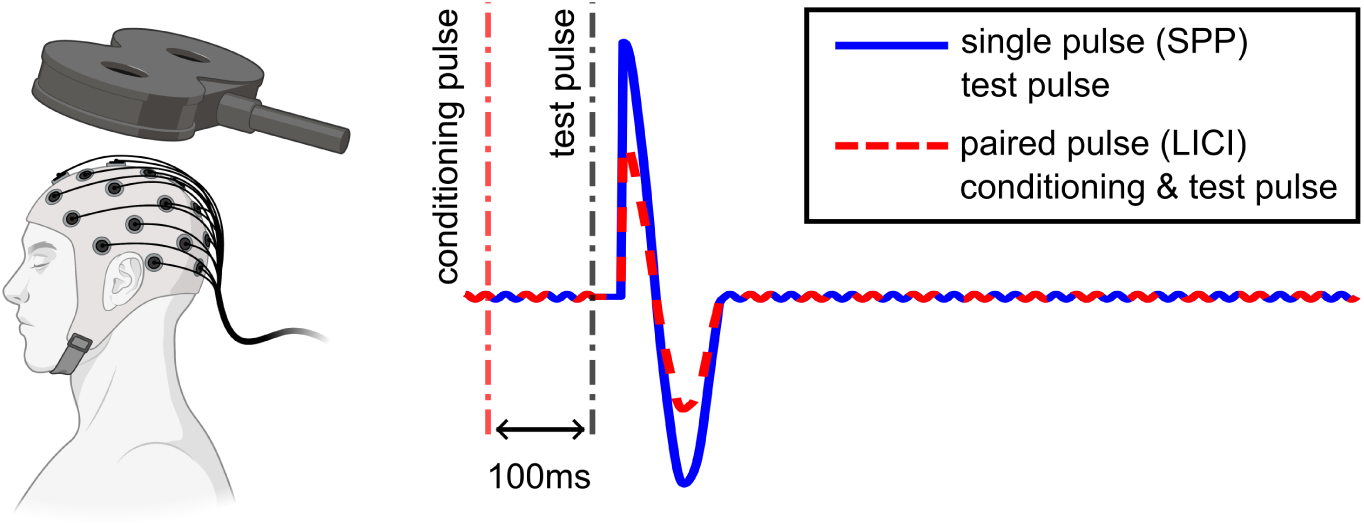
Schematic illustration of the Long Interval intracortical inhibition (LICI) paired-pulse paradigm. (Left) Transcranial Magnetic Stimulation (TMS) applied over the frontal cortex during EEG recording. (Right) Stimulation timing with a 100 ms inter-stimulus interval (ISI) separating the conditioning pulse (*t* = *−*100 ms) and the test pulse (*t* = 0 ms). The traces compare a single pulse (SPP, solid blue line) with the paired-pulse condition (LICI, dashed red line), depicting cortical suppression induced by the conditioning stimulus. Waveforms are schematic representations intended for conceptual illustration only and do not reflect empirical recorded data.

### 2.4. Data acquisition and analysis

To convert the multivariate raw EEG time series data into discrete metrics evaluating the overall brain dynamics following single or paired-pulse TMS, dynamical metrics were extracted for each participant from the averaged EEG data across the following domains: 1. Temporal features based on 20 channels across six time blocks (50–200, 200–350, 350–500, 500–650, 650–800, 800–950 ms), 2. Frequency features across the five frequency bands (delta, theta, alpha, beta, and gamma) using the entire EEG trace (50–950 ms), and 3. Time-frequency features extracted over the six time blocks and five frequency bands. Described further in Figure 2.

**Figure 2:**
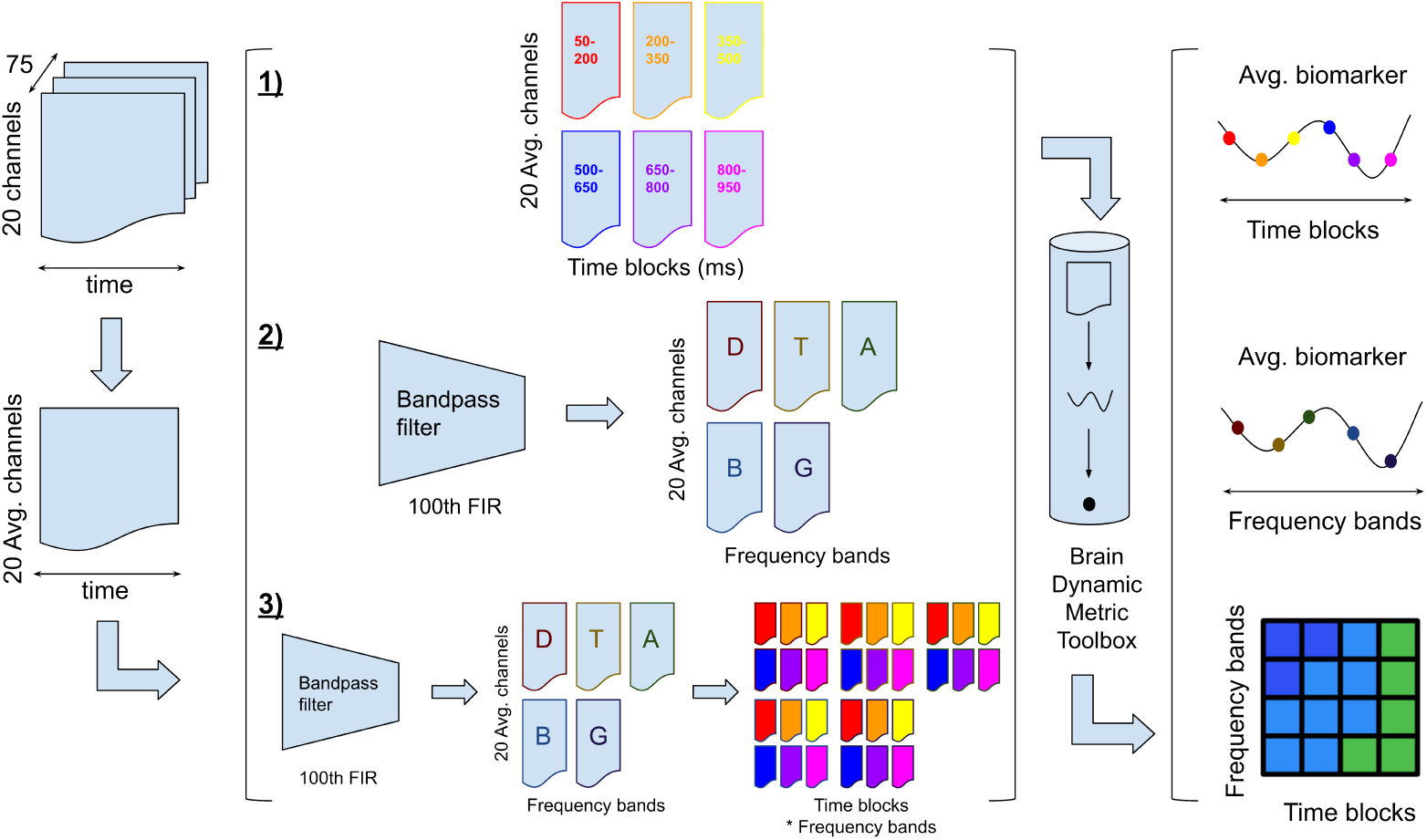
Data Preparation Pipeline: The dataset comprises recordings from 20 channels EEG across 75 trials per stimulation site and condition, spanning 50 to 950 ms post-test-stimulus for each participant. Processing involves three distinct analyses focusing on temporal, frequency, and temporal frequency dimensions. For spectral and time frequency analyses, continuous 50 to 950 ms epochs are first bandpass filtered into five canonical frequency bands (Delta, Theta, Alpha, Beta, and Gamma) using a 100th order Finite Impulse Response (FIR) filter prior to temporal segmentation into six 150 ms blocks. Dynamical metrics are extracted across the 20 channels using the Brain Dynamic Metric Toolbox (https://github.com/qinwayne/BrainDynamicMetricToolbox) and averaged across valid trials to yield a regional summary metric for each participant, condition, temporal block, and frequency band.

#### 2.4.1. Dynamical metrics

Perturbative Integration Latency Index (PILI) is a perturbation-based metric that quantifies the residual energy injected into a system by a perturbation over time. It reflects the cortical excitability and resilience of the brain to external disturbances. PILI is calculated as the area under the curve (signal), which indicates how long it takes for the system to return to its normal state. The recovery rate of the system depends on the time constant *τ*, which determines how fast the energy dissipates. The PILI can be computed by computing the area under the curve of the signal multiplied by its sampling frequency to estimate the energy of the signal. The larger the area, the more energy is present in the signal. Moreover, the change of the area can reflect the rate of energy dissipation from the system. It can provide information about the degree of integration and latency of the brain region.

The autocorrelation function (ACF) is a measure of how similar a signal is to itself at different time lags. It can be used to analyse the temporal properties and independence of signals, such as electroencephalogram (EEG) signals used in this study. The ACF width of EEG signals reflects the degree of correlation between adjacent time segments, which implies the degree of brain activity or synchronisation. A widely adopted way to analyse the time series data is to apply a moving window technique that splits the data into smaller segments. Then, for each segment, the autocorrelation function (Matlab xcorr) can be calculated and its width can be determined by finding the point where it drops to 34.1% of its maximum value. A high ACF width indicates a high degree of brain activity or synchronisation, while a low ACF width indicates a low degree of brain activity or synchronisation. Autocorrelation can be used to study the temporal structure and variability of EEG signals.

Lyapunov exponents (LE) are measures of the divergence and convergence rate of a dynamical system. They indicate how fast two closely embedded trajectories in phase space separate from each other over time, rather than focusing on local behaviours near a fixed point. LE can be estimated using time-series data and are useful for determining the stability of oscillations. Each dimension of the phase plane has a corresponding LE that shows the rate of expansion or contraction of orbits in that dimension, with positive LE indicating local instability and negative LE indicating local stability. The distance between two points on two close trajectories in phase space can be expressed as *|δ*(*t*)*|* = *|δ*(0)*|e^λt^*, where *δ*(0) is the initial distance and *λ* is the LE, assuming an exponential rate of divergence. To implement LE, it is required to first estimate the dimension of the system. A suitable embedding dimension can be chosen by using Cao’s embedding algorithm (Cao, 1997). In a defined high-dimensional space, the Wolf method (as shown in Table 1) tracks the trajectories of two points as they evolve and calculates the LE based on the length of the principal axis of the elliptical orbits that they follow. The overall stability of a system is determined by the sum of the LE. In this study, we used lag-1 embedding approach to embed the data into m dimensions from which to estimate the LE metric from the EEG data.

**Table 1:**
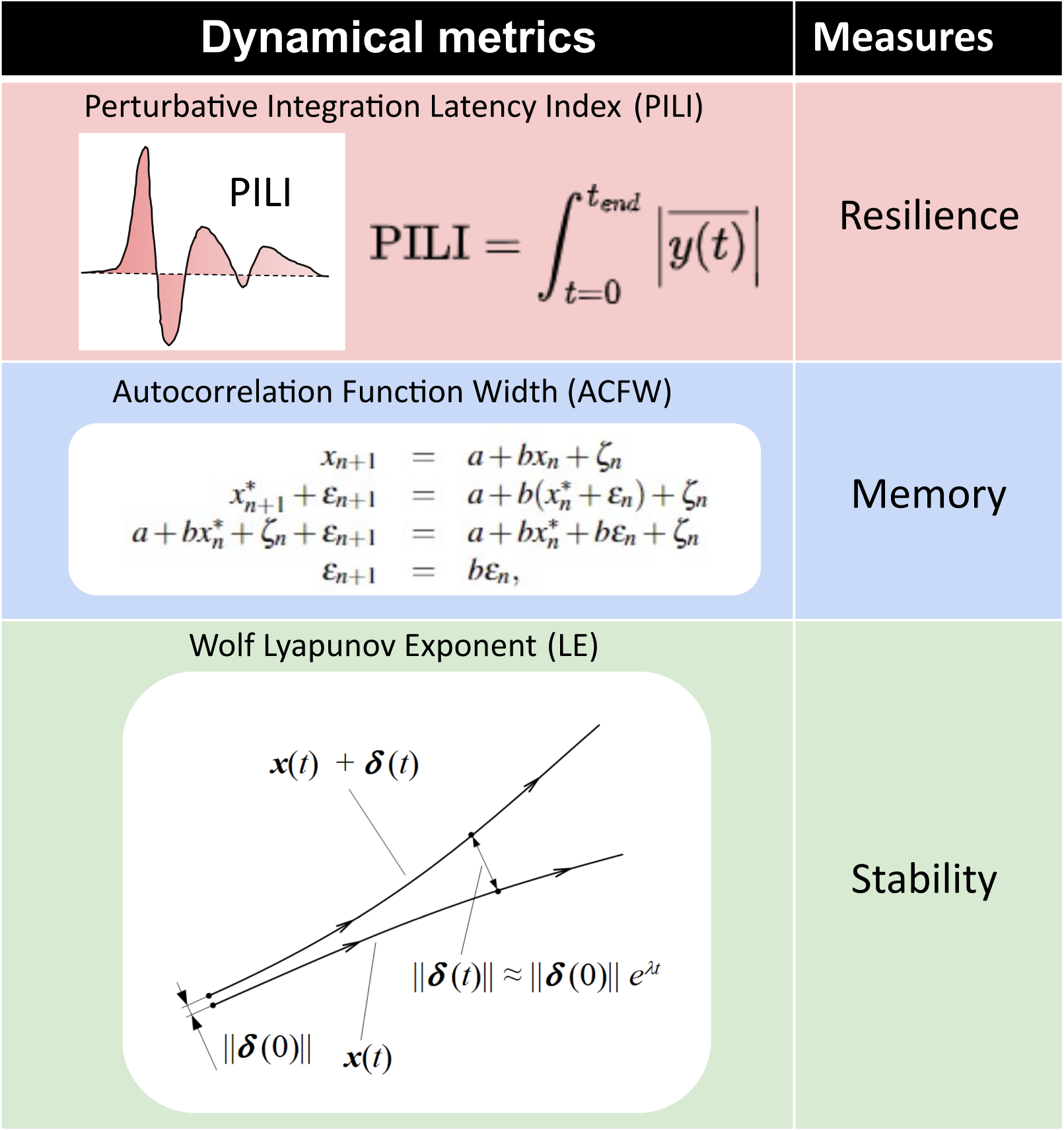
Overview of the three distinct dynamical metrics utilised to analyse the time-series data: Perturbative Integration Latency Index (PILI), Autocorrelation Function Width (ACFW), and Lyapunov Exponent (LE). The PILI quantifies the system’s recovery and resilience following Transcranial Magnetic Stimulation (TMS) by calculating the area under the curve of the baseline-corrected response Jobst et al. (2017). The ACFW assesses the temporal memory and signal interdependencies by measuring the thresholded decay of the autocorrelation function Maturana et al. (2020). Lastly, the LE evaluates the chaotic dynamics and oscillatory stability of the neural system by leveraging Wolf’s algorithm (Wolf et al., 1985) for phase-space reconstruction to measure the exponential divergence of trajectories.

| Dynamical metrics | Measures |
|---|---|
| Perturbative Integration Latency Index (PILI)<br>PILI<br>$\mathrm{PILI} = \int_{t=0}^{t_{end}} \left\|\overline{y(t)}\right\|$ | Resilience |
| Autocorrelation Function Width (ACFW)<br>x<sub>n+1</sub> = a + bx<sub>n</sub> + ζ<sub>n</sub><br>x<sup>*</sup><sub>n+1</sub> + ε<sub>n+1</sub> = a + b(x<sup>*</sup><sub>n</sub> + ε<sub>n</sub>) + ζ<sub>n</sub><br>a + bx<sup>*</sup><sub>n</sub> + ζ<sub>n</sub> + ε<sub>n+1</sub> = a + bx<sup>*</sup><sub>n</sub> + bε<sub>n</sub> + ζ<sub>n</sub><br>ε<sub>n+1</sub> = bε<sub>n</sub>, | Memory |
| Wolf Lyapunov Exponent (LE)<br>**x**(t) + **δ**(t)<br>\|\|**δ**(t)\|\| ≈ \|\|**δ**(0)\|\| e<sup>λt</sup><br>\|\|**δ**(0)\|\|<br>**x**(t) | Stability |

### 2.5. Brain Dynamic Metric Toolbox

From this study, authors have developed the Brain Dynamic Metric Toolbox which has been created as an open-source and user-friendly Github repository (https://github.com/qinwayne/BrainDynamicMetricToolbox). While this toolbox was developed within the context of the described TMS case study, it can be adapted to other modalities of brain stimulation when an estimation of brain dynamics is required. All documentation, data requirements, and relevant codes relating to the Brain Dynamic Metric Toolbox are contained within the Github repository.

### 2.6. Statistical models

In this investigation, we employed a Linear Mixed-Effects (LME) model in R, using the lme4 package (Bates et al., 2015), to perform statistical analysis on results. The dependent variable was the TMS-evoked potential following either single or paired-pulse TMS. A linear mixed model was fitted using restricted maximum likelihood (REML). The model explored various metrics over time (temporal features over 6 time blocks) and/or frequency (spectral features based on 5 frequency blocks), side (left or right stimulation), and pulse (single or paired-pulse). The initial analysis sought to determine which combination of these variables resulted in the most parsimonious model, where the full model was given by Equation 1

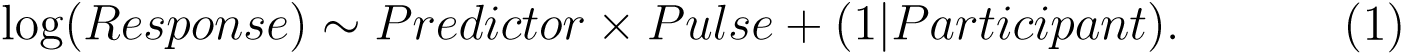

In this equation, “*Response*” represents the feature of interest, and “*Predictor*” is the explanatory variable, which is one of “*Time*”, “*Frequency*”, or “*Temporal − Frequency*”. “*Pulse*” refers to the two TMS paradigms (single or paired). “*Participant*” was incorporated as a random effect in the model, represented by the term “1*|Participant*”. Side indicates the location of the TMS applied. A model comparison analysis was performed, and it was found that excluding the “*Side*” factor achieved a lower AIC value (1513.43) compared to the model that included it (AIC = 1533.66), therefore removing side resulted in a better description of the relationship between the predictors and the dependent variables.

The effect of the type of pulse (single versus paired) for each dynamical metric (PILI, LE, and ACWF) was explored across these temporal and spectral domains. To account for multiple comparisons, Bonferroni’s correction was applied. Response variables, which were initially log-transformed, were back-transformed following the post-hoc analysis to facilitate the interpretation and reporting of the findings. To address negative raw LE values, transformations (log and sqrt) were evaluated as absolute values. The post-hoc analysis was conducted to identify significant contrasts with respect to the explanatory variables compared across pulse types (single vs paired-pulse) for each of the perturbational metrics. For each of the temporal, spectral, and tempo-frequency studies, we generated plots to display the general trend changes of the metrics (PILI, ACFW, and LE) across different blocks. To ensure the integrity of the analysis, cases with missing observed values were excluded, maintaining a dataset of complete cases only.

## 3. Results

### 3.1. Temporal features

PILI quantifies the cortical resilience following TMS perturbation. The LME model included participants as a random effect and showed substantial explanatory power (conditional R^2^ of 0.70), indicating that 70% of the variation in the data was accounted for by the model, with fixed effects (marginal R^2^ of 0.66) representing 66% of the variation. Analysis of the fixed effects described in Table 2 revealed significant effects of time (F(5,319.4) = 152, p *<* 0.001) and the interaction between time and pulse (F(5,319.4) = 3, p = 0.01). Post-hoc analysis was conducted to further identify significant contrasts with respect to pulse type (comparing single and paired-pulse) across the six time-points. A significant difference between pulse type was identified at the first timepoint (150 ms) with paired-pulse showing inhibition (EM = 0.042, SD = 0.0116) relative to the single pulse (EM = 0.050, SD = 0.0138). From 300 ms onwards, PILI values for both pulse types converged with no significant differences between them, indicating similar levels of residual energy as they returned to baseline (Figure 3A).

**Figure 3:**
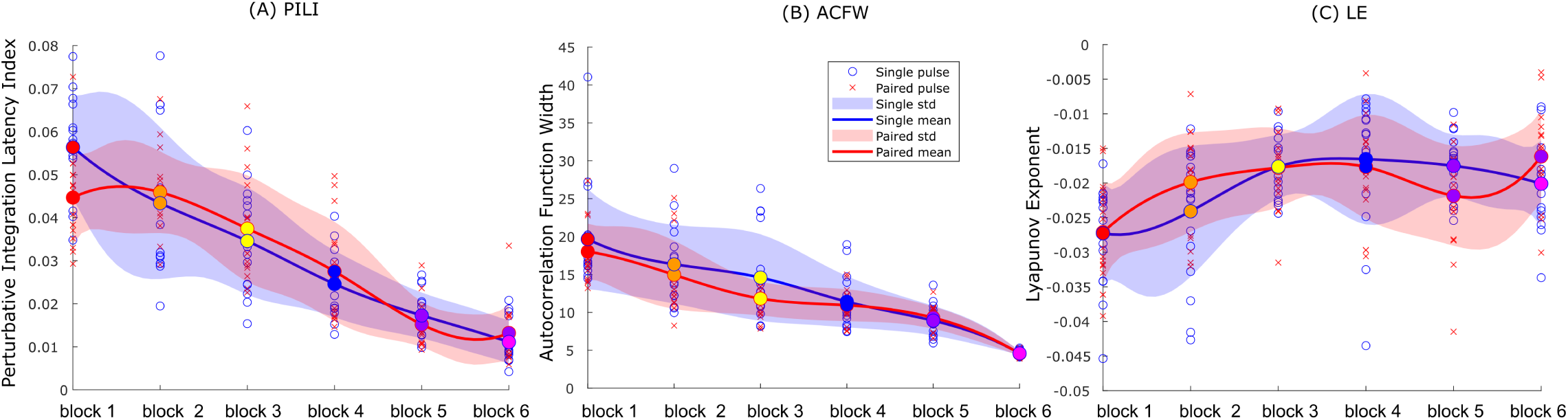
Temporal features: Dynamical metrics reveal the temporal alterations induced by Transcranial Magnetic Stimulation (TMS). The horizontal axis denotes consecutive post-stimulation time windows: block 1 (150 ms), block 2 (300 ms), block 3 (450 ms), block 4 (600 ms), block 5 (750 ms), and block 6 (900 ms). Round dots represent the averaged data per participant for each condition, where blue dots denote single-pulse TMS and red dots denote paired-pulse TMS. Shaded regions indicate standard deviation. 3A. Perturbative Integration Latency Index (PILI) discerns initial disparities when contrasting single-pulse and paired-pulse TMS protocols. 3B. Autocorrelation Function Width (ACFW) exhibits a diminishing trend over time subsequent to TMS perturbation. Concurrently, 3C. the Lyapunov Exponent (LE) gradually moves towards 0. Note that the data is not continuous but is presented this way for illustrative purposes.

**Table 2:** Temporal features: Linear mixed model (LMM) reveals the interactions between the dynamical metrics and the temporal alterations induced by Transcranial Magnetic Stimulation (TMS).

|  | **PILI** |  |  |  | **ACFW** |  |  |  | **LE** |  |  |  |
|---|---|---|---|---|---|---|---|---|---|---|---|---|
| **term** | **df1** | **df2** | **F.ratio** | **p.value** | **df1** | **df2** | **F.ratio** | **p.value** | **df1** | **df2** | **F.ratio** | **p.value** |
| *time* | 5 | 319.4 | 152 | <**0.001** | 5 | 319.1 | 425.9 | <**0.001** | 5 | 319.7 | 15.2 | <**0.001** |
| *pulse* | 1 | 331.6 | 0.62 | 0.43 | 1 | 325.5 | 13.84 | <**0.001** | 1 | 332.9 | 1.02 | 0.31 |
| *time:pulse* | 5 | 319.4 | 3 | **0.01** | 5 | 319.1 | 2.93 | **0.01** | 5 | 319.7 | 3.38 | <**0.001** |

*Note.* PILI = Perturbative Integration Latency Index; ACFW = Autocorrelation Function Width; LE = Lyapunov Exponent. Bold values indicate statistical significance.

ACFW quantifies the self-similarity or temporal memory of brain activity following TMS perturbation. The LME model demonstrated substantial explanatory power (conditional R-squared of 0.87) with marginal R-squared of 0.80. Analysis of fixed effects, as described in Table 2, revealed significant main effects of time (F(5,319.1) = 425.9, p *<* 0.001), pulse type (F(1,325.5) = 13.84, p *<* 0.001) and time*pulse type (F(5,319.1) = 2.93, p = 0.01). Post-hoc analysis identified a significant difference between pulse types in the early time intervals (150 to 450 ms), with paired-pulse stimulation exhibiting inhibition (EM = 18.26, SD = 4.33) relative to single-pulse stimulation (EM = 20.38, SD = 6.28) at 150 ms, paired-pulse (EM = 14.51, SD = 4.02) vs single-pulse (EM = 16.32, SD = 4.80) at 300 ms, paired-pulse (EM = 11.73, SD = 2.15) vs single-pulse (EM = 14.90, SD = 5.89) at 450 ms. From 450 ms onwards, ACFW values for both pulse types converged, indicating similar levels of temporal memory as they returned to baseline (Figure 3B).

LE quantifies the phase space trajectory stability of oscillatory brain responses following TMS perturbation. The LME model exhibited low explanatory power (conditional R-squared of 0.26), with marginal R-squared of 0.20, suggesting there are likely other causes of variability. Analysis of the fixed effects, as described in Table 2, revealed significant main effects of time (F(5,319.7) = 15.2, p *<* 0.001) and a significant interaction between time and pulse type (F(5,319.7) = 3.38, p *<* 0.001). Post-hoc analysis was also conducted to identify significant contrasts between single-pulse and paired-pulse stimulation across the six time-points. A significant difference between pulse types was observed at the 300 ms timepoint and the 900 ms timepoint, with paired-pulse stimulation exhibiting variations (300 ms: EM =-0.020, SD = 0.006; 900 ms: EM =-0.017, SD = 0.006; Bonferroni-adjusted contrast on log-transformed magnitudes, p *<* 0.05) relative to single-pulse stimulation (300 ms: EM =-0.025, SD = 0.009; 900 ms: EM =-0.019, SD = 0.006). At other timepoints, the responses from the two paradigms converged, as shown in Figure 3C.

### 3.2. Frequency features

Following this, the frequency domain features were explored across the three perturbational metrics. For this analysis, a square root transformation was required to ensure the data conformed to a normal distribution. The results are presented in Table 3.

**Table 3:** Linear mixed model (LMM) reveals the interactions between the dynamical metrics and the spectrum alterations induced by Transcranial Magnetic Stimulation (TMS). Results from the LMM match what is seen in Figure 4.

|  | **PILI** |  |  |  | **ACFW** |  |  |  | **LE** |  |  |  |
|---|---|---|---|---|---|---|---|---|---|---|---|---|
| **term** | **df1** | **df2** | **F.ratio** | **p.value** | **df1** | **df2** | **F.ratio** | **p.value** | **df1** | **df2** | **F.ratio** | **p.value** |
| *frequency* | 4 | 263.3 | 760.1 | <**0.001** | 4 | 263.2 | 1293 | <**0.001** | 4 | 263.4 | 209.1 | <**0.001** |
| *pulse* | 1 | 273.7 | 1.91 | 0.17 | 1 | 272 | 5.4 | **0.02** | 1 | 273.9 | 7.5 | <**0.01** |
| *frequency:pulse* | 4 | 263.3 | 0.15 | 0.96 | 4 | 263.2 | 5.82 | <**0.001** | 4 | 263.4 | 1.55 | 0.19 |

*Note.* PILI = Perturbative Integration Latency Index; ACFW = Autocorrelation Function Width; LE = Lyapunov Exponent. Bold values indicate statistical significance.

For the PILI metric measured responses, the LME model, showed good explanatory power (conditional R-squared of 0.91), with marginal R-squared of 0.90. Analysis of the fixed effects, presented in Table 3, revealed significant effects of frequency ranges (F(4,263.3) = 760.1, p *<* 0.001), but no significant interactions (Figure 4A).

**Figure 4:**
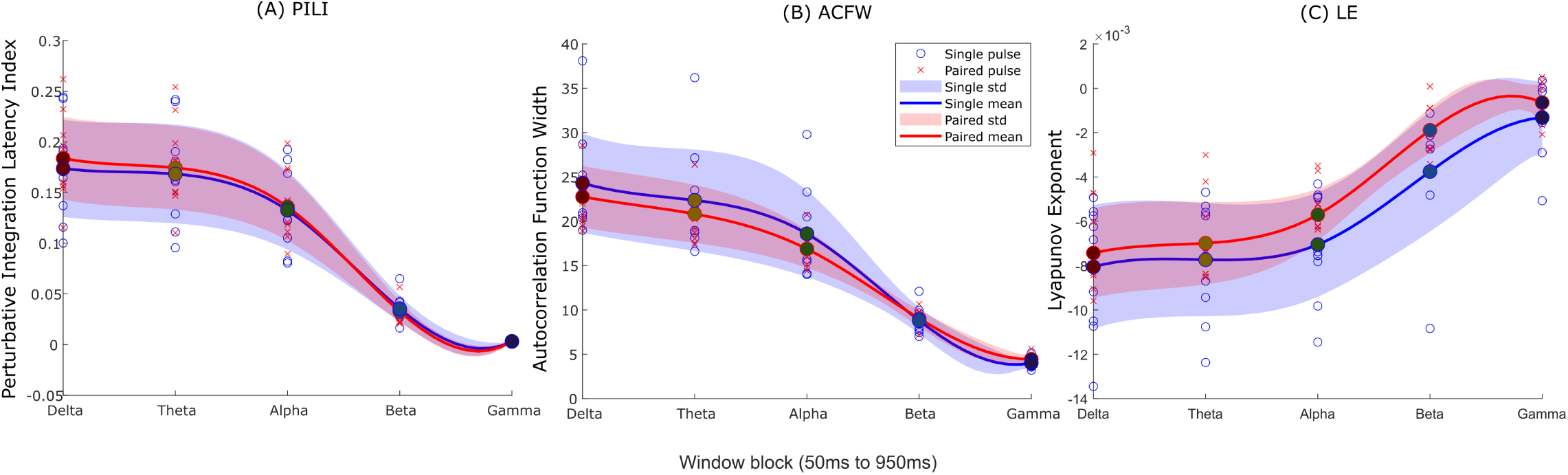
Spectral features: Dynamical metrics reveal the spectrum alterations induced by Transcranial Magnetic Stimulation (TMS). 4A. Perturbative Integration Latency Index (PILI) and the 4B. Autocorrelation Function Width (ACFW) exhibit high similarity between two paradigms, with the delta band containing the most information and the gamma band the least. Concurrently, 4C. the Lyapunov Exponent (LE) gradually approaches zero. Note that the data is not continuous but is presented this way for illustrative purposes.

For the ACFW metric, the LME model showed substantial explanatory power (conditional R-squared of 0.95), with marginal R-squared of 0.93. Analysis of the fixed effects, as described in Table 3, revealed significant effects of frequency ranges (F(4,263.2) = 1293, p *<* 0.001), pulse type (F(1,272) = 5.4, p = 0.02), and frequency*pulse type (F(4,263.2) = 5.82, p *<* 0.001). Post-hoc analysis revealed significant differences between pulse types across all frequency bands except for Beta. The ACFW value trends for single-pulse and paired-pulse stimulation were similar, but responses from the paired-pulse paradigm were lower than those from the single-pulse paradigm in the low-frequency domains (Delta-EM: 22.71 vs. 24.79, SD: 3.24 vs. 4.20, Theta-EM: 20.36 vs. 22.52, SD: 3.09 vs. 4.31, and Alpha-EM: 16.19 vs. 18.19, SD: 2.20 vs. 4.01) and higher in the high-frequency Gamma domain (EM: 4.37 vs. 4.12, SD: 0.41 vs. 0.46) (Figure 4B).

For the LE metric, the LME model also presented with substantial explanatory power (conditional R-squared of 0.76), with marginal R-squared of 0.72. Analysis of the fixed effects, as described in Table 3, revealed significant effects of frequency range (F(4,263.4) = 209.1, p *<* 0.001) and pulse type (F(1,273.9) = 7.5, p *<* 0.01), indicating slightly less negative LE values overall during paired pulse relative to single pulse stimulation across the spectrum, but no significant interactions (Figure 4C).

### 3.3. Temporal * Frequency

For the temporal*frequency analysis, we evaluated response alterations by calculating the direct ratio of paired pulse stimulation to single pulse stimulation 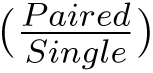 across each frequency band and temporal window. The results are presented in Table 4. Under this scale, a ratio of 1.0 indicates parity between stimulation protocols as shown in Figure 5. Values above 1.0 signify paired pulse predominance (shown in red), whereas values below 1.0 indicate cortical suppression where single pulse responses exceed paired pulse responses (shown in blue). Values were averaged across both left and right hemispheric stimulation blocks. Post hoc contrasts were evaluated to identify the temporal and spectral blocks displaying significant differences between pulse types, with significant cells denoted by asterisks.

**Figure 5:**
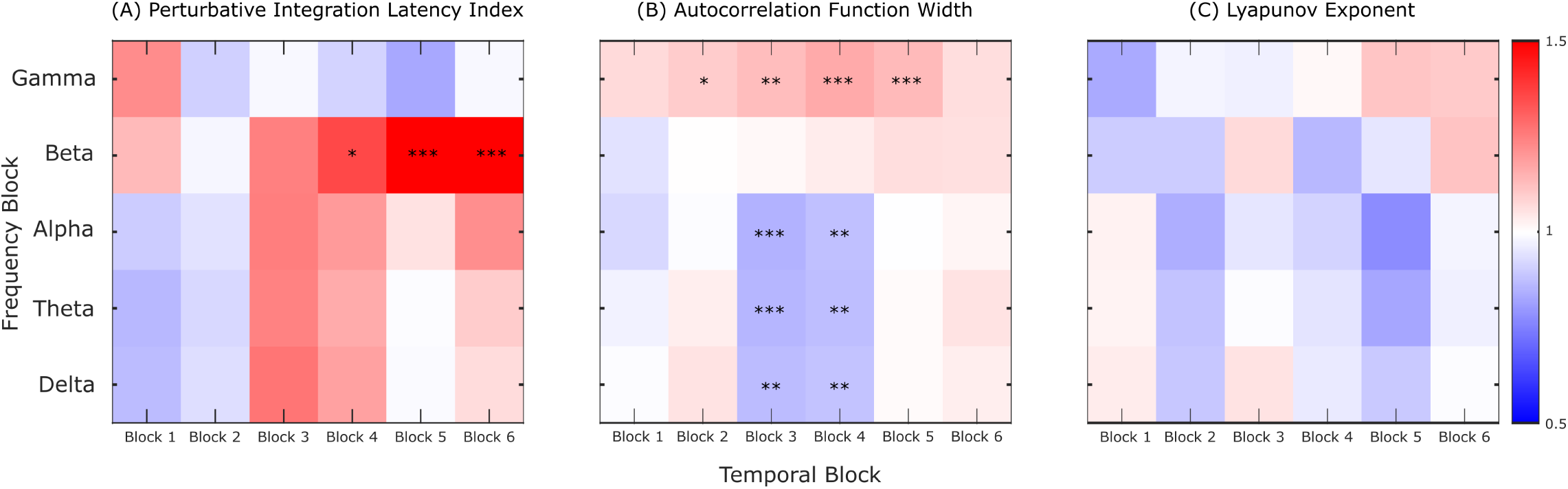
Temporal-Frequency features: Ratio comparison (paired/single pulse) of dynamical metrics reveals the temporal-spectral alterations induced by Transcranial Magnetic Stimulation (TMS). Colour scale represents the direct ratio of paired pulse to single pulse responses bounded between 0.5 and 1.5, with 1.0 indicating equal response magnitudes. 5A. Perturbative Integration Latency Index (PILI) shows significant differences exclusively in the beta band across later time windows (Block 4 to Block 6, corresponding to 600 ms to 900 ms). Asterisks denote statistically significant post hoc differences (* p *<* 0.05, ** p *<* 0.01, *** p *<* 0.001). 5B. Autocorrelation Function Width (ACFW) exhibits significant differences across low frequency bands (Delta, Theta, Alpha) in Block 3 and Block 4 (450 ms to 600 ms), alongside sustained differences in the gamma band from Block 2 to Block 5 (300 ms to 750 ms). In contrast, 5C. the Lyapunov Exponent (LE) displays no statistically significant differences across any temporal or spectral blocks. Asterisks denote significant post hoc contrasts.

**Table 4:** Linear mixed model (LMM) analysis detailing the statistical interactions between dynamical metrics and the temporal-frequency alterations induced by single-and paired-pulse Transcranial Magnetic Stimulation (TMS). The LMM results quantitatively support the visual trends presented in Figure 5.

|  | **PILI** |  |  |  | **ACFW** |  |  |  | **LE** |  |  |  |
|---|---|---|---|---|---|---|---|---|---|---|---|---|
| **term** | **df1** | **df2** | **F.ratio** | **p.value** | **df1** | **df2** | **F.ratio** | **p.value** | **df1** | **df2** | **F.ratio** | **p.value** |
| *time* | 5 | 1663 | 90.71 | <**0.001** | 5 | 1663 | 3730 | <**0.001** | 5 | 1664 | 17.4 | <**0.001** |
| *frequency* | 4 | 1663 | 4413 | <**0.001** | 4 | 1663 | 7382 | <**0.001** | 4 | 1664 | 356.1 | <**0.001** |
| *pulse* | 1 | 1662 | 19.82 | <**0.001** | 1 | 1679 | 0.56 | 0.46 | 1 | 1605 | 3.64 | 0.06 |
| *time:freq.* | 20 | 1663 | 117.2 | <**0.001** | 20 | 1663 | 287.9 | <**0.001** | 20 | 1664 | 1.74 | **0.02** |
| *time:pulse* | 5 | 1663 | 4.23 | <**0.001** | 5 | 1663 | 10.33 | <**0.001** | 5 | 1664 | 0.85 | 0.52 |
| *freq.:pulse* | 4 | 1663 | 11.82 | <**0.001** | 4 | 1663 | 18.58 | <**0.001** | 4 | 1664 | 0.46 | 0.76 |
| *t.:freq.:pul.* | 20 | 1663 | 1.73 | **0.02** | 20 | 1663 | 2.19 | <**0.01** | 20 | 1664 | 0.97 | 0.5 |

*Note.* PILI = Perturbative Integration Latency Index; ACFW = Autocorrelation Function Width; LE = Lyapunov Exponent; df = degrees of freedom. Bold values indicate statistical significance.

For the PILI metric, the LME model showed substantial explanatory power (conditional R-squared of 0.92), with marginal R-squared of 0.92. Analysis of the fixed effects, as described in Table 4, revealed significant effects of time (F(5,1663) = 90.71, p *<* 0.001), frequency (F(4,1663) = 4413, p *<* 0.001), pulse (F(1,1662) = 19.82, p *<* 0.001), and time*frequency (F(20,1663) = 117.2, p *<* 0.001), time*pulse (F(5,1663) = 4.23, p *<* 0.001), frequency*pulse (F(4,1663) = 11.82, p *<* 0.001), as well as time*frequency*pulse (F(20,1663) = 1.73, p = 0.02). Post-hoc analysis revealed three significant differences between pulse types with data described in Table 5 and Figure 5A.

**Table 5:** PILI Temporal Frequency Post-Hoc Analysis Summary.

| **Time** | **Frequency** | **EM<sub>paired</sub>** | **EM<sub>single</sub>** | **SD<sub>paired</sub>** | **SD<sub>single</sub>** |
|---|---|---|---|---|---|
| 600 | Beta | 0.012 | 0.0088 | 0.0040 | 0.0029 |
| 750 | Beta | 0.013 | 0.0088 | 0.0034 | 0.0042 |
| 900 | Beta | 0.018 | 0.010 | 0.0064 | 0.0052 |

*Note.* EM = Estimated Marginal Means; SD = Standard Deviation. Data represents paired-pulse and single-pulse Transcranial Magnetic Stimulation (TMS) across time points (ms). Values are rounded to two significant digits.

For the ACFW metric, the LME model had strong explanatory power (conditional R-squared of 0.97), with marginal R-squared of 0.97. Analysis of the fixed effects, as described in Table 4, revealed significant effects of time (F(5,1663) = 3730, p *<* 0.001), frequency (F(4,1663) = 7382, p *<* 0.001), and time*frequency (F(20,1663) = 287.9, p *<* 0.001), time*pulse (F(5,1663) = 10.33, p *<* 0.001), frequency*pulse (F(4,1663) = 18.58, p *<* 0.001), and time*frequency*pulse (F(20,1663) = 2.19, p *<* 0.01). Post-hoc analysis revealed several significant differences between pulse types with data described in Table 6 and Figure 5B.

**Table 6:** ACFW Temporal Frequency Post-Hoc Analysis Summary.

| **Time** | **Frequency** | **EM<sub>paired</sub>** | **EM<sub>single</sub>** | **SD<sub>paired</sub>** | **SD<sub>single</sub>** |
|---|---|---|---|---|---|
| 450 | Delta | 26 | 30 | 5.3 | 3.4 |
| 600 | Delta | 22 | 24 | 3.3 | 4.9 |
| 450 | Theta | 23 | 27 | 5.4 | 3.5 |
| 600 | Theta | 20 | 22 | 3.6 | 5.2 |
| 450 | Alpha | 18 | 21 | 4.9 | 3.7 |
| 600 | Alpha | 16 | 18 | 3.3 | 4.9 |
| 300 | Gamma | 4.6 | 4.2 | 0.31 | 0.37 |
| 450 | Gamma | 4.7 | 4.2 | 0.31 | 0.47 |
| 600 | Gamma | 4.7 | 4.1 | 0.31 | 0.44 |
| 750 | Gamma | 4.5 | 4.0 | 0.51 | 0.42 |

*Note.* EM = Estimated Marginal Means; SD = Standard Deviation. Data represents paired-pulse and single-pulse Transcranial Magnetic Stimulation (TMS) across time points (ms) for the Autocorrelation Function Width (ACFW). Values are rounded to two significant digits.

For the LE metric, the LME model showed low explanatory power (conditional R-squared of 0.45), with marginal R-squared of 0.45. Given the low explanatory power, it is important to note that there are likely other causes of variability. Analysis of the fixed effects, as described in Table 4, revealed significant effects of time (F(5,1664) = 17.4, p *<* 0.001), frequency (F(4,1664) = 356.1, p *<* 0.001), and significant interactions between time and frequency (F(20,1664) = 1.74, p = 0.02). Post-hoc analysis revealed no significant differences between pulse types as seen in Figure 5C.

## 4. Discussion

In the present study, we applied a perturbation–recovery framework to frontal TMS-EEG responses evoked by single-pulse stimulation and paired-pulse long-interval intracortical inhibition (LICI). The principal finding was that paired-pulse stimulation altered the temporal and spectral profile of the evoked cortical response, particularly when quantified using metrics sensitive to cortical resilience and temporal memory. Across analyses, the Perturbative Integration Latency Index (PILI) and Autocorrelation Function Width (ACFW) were the most consistent measures for distinguishing single-pulse and paired-pulse responses. The Lyapunov Exponent (LE) provided an additional estimate of post-stimulation stability but showed weaker and less consistent discrimination between stimulation conditions. These findings support the value of treating TMS-EEG responses as spatiotemporal dynamics rather than only quantifying amplitude response at selected latencies. In the context of LICI, the effect of the conditioning pulse was expressed not only as suppression of the evoked response, but also as altered resilience and temporal memory of the post-stimulation signal. This suggests that dynamical metrics may provide complementary information about inhibitory physiology that is not captured by conventional amplitude-based summaries alone.

### 4.1. Paired-pulse inhibition alters perturbation-recovery dynamics

LICI is commonly interpreted as reflecting cortical inhibitory mechanisms, including processes consistent with GABA_B_ receptor-mediated inhibition (McDonnell et al., 2006; Daskalakis et al., 2008; Rogasch et al., 2013; Fatih et al., 2021). In the present study, paired-pulse stimulation was associated with altered cortical resilience and reduced temporal memory, as indexed by PILI and ACFW. This pattern is consistent with the expected suppression effect of the conditioning pulse but suggests that its influence extends beyond a reduction in evoked-response magnitude (peak amplitude).

A key implication is that inhibitory conditioning may alter how the cortical response evolves after stimulation. If the conditioning pulse constrains the response to the test pulse, this may be expressed not only as a smaller initial response, but also as less temporally persistent response trajectory. In this framework, inhibition is not only reflected in whether a response is smaller at a given latency or electrode, but also in the temporal dynamics of the response: how long stimulation-related activity remains structured before dissipating towards baseline. This interpretation is particularly relevant for clinical populations in which excitation-inhibition balance is thought to be altered, because abnormal inhibition may be expressed through changes in response dynamics as well as response magnitude. The interpretation of LICI as reflecting GABA_B_-mediated inhibition is supported by pharmacological studies demonstrating that the GABA_B_ receptor agonist baclofen enhances LICI, including when responses are measured directly using TMS-EEG (McDonnell et al., 2006; Premoli et al., 2014). However, PILI and ACFW are electrophysiological measures derived from scalp EEG and while their modulation is compatible with the expected effects of LICI, they do not directly measure synaptic inhibition or GABA_B_-receptor function.

Against this mechanistic background, the present findings extend previous frontal TMS-EEG research on cortical inhibition in neuropsychiatric populations. Prior studies have characterised LICI through suppression of evoked cortical response and oscillatory activity in frontal and motor regions across various psychiatric disorders, such as schizophrenia and alcohol dependence (Farzan et al., 2010; Radhu et al., 2013; Naim-Feil et al., 2016; Fatih et al., 2021). More recent work has extended this perturbational approach to the organisation of distributed network responses following inhibitory stimulation (Naim-Feil et al., 2022). The present study builds on this foundation by examining an additional dimension of the response, its temporal dynamics, resilience and recovery, following a perturbation. These findings therefore provide a complementary perspective on LICI, in which the effect of inhibition is characterised not only by how strongly the cortex responds, but also by how that response unfolds over time.

### 4.2. Temporal and spectral dynamics

The temporal findings highlight the importance of analysing the broader post-stimulation response beyond 300 ms. While conventional TMS-EEG analyses generally focus on discrete evoked components within the first several hundred milliseconds following stimulation, they may not fully describe how the evoked response persists, decays, or changes in temporal structure after stimulation. By extending the analysis to 950 ms post-stimulation, the present framework allowed single-pulse and paired-pulse responses to be compared as a dynamical response across a longer post-perturbation trajectory (Pigorini et al., 2011; Mutanen et al., 2013).

This approach builds on previous work conceptualising the TMS-evoked response as a dynamic perturbation rather than a series of discrete evoked components. Mutanen et al. (2013), for example, used recurrence analysis to track state-space trajectories of EEG activity before and after single-pulse TMS. They demonstrate that TMS alters the state-space trajectory and temporal evolution of brain state. Subsequent TMS–EEG and connectome studies have shown that the propagation of evoked activity is shaped by the structural organisation of the stimulated network (Momi et al., 2021).

Importantly, in our dataset, the effect of inhibitory conditioning was not temporally uniform and differed across dynamical measures. The early reduction in PILI suggests that the most prominent effect on cortical resilience and residual energy occurred relatively soon after stimulation, with single-and paired-pulse trajectories subsequently converging. However, changes in ACFW persisted for longer, suggesting that inhibitory conditioning continued to alter the temporal memory of cortical activity after differences in overall response energy had diminished. This indicates that response magnitude and temporal organisation may capture distinct, and potentially more relevant, features of the system’s dynamic and nonlinear response to inhibitory conditioning. The time-frequency analysis further showed that later differences were frequency-specific rather than reflecting sustained global suppression, with effects emerging across different temporal windows in beta, lower-frequency, and gamma activity. Together, these findings support the central premise of the present framework: that paired-pulse inhibition modulates a range of dynamical properties over the evolving cortical response rather than as a uniform reduction in response amplitude.

More recent in-silico network lesion experiments provide a mechanistic framework for interpreting these temporal dynamics. Using subject-specific connectome-based models of TMS-EEG responses, Momi et al. (2023) showed that early components can arise largely from local dynamics, whereas later components increasingly depend on recurrent interactions across distributed brain networks (Momi et al., 2023). Subsequent modelling with direct intracerebral electrical stimulation suggests that the contribution of recurrent connectivity to later evoked responses varies across cortical networks and may be particularly important in higher-order association networks (Momi et al., 2025). Convergent evidence from studies combining TMS with intracranial recordings further demonstrates that the effects of stimulation extend beyond the directly stimulated region to remote cortical sites and broader network interactions (Tsang et al., 2026). Together, these findings support the view that the evolving post-stimulation activity reflects interactions within a distributed dynamical system, rather than a sequence of isolated local responses.

The present study examines a complementary dimension of this process. Whereas computational and connectome-based approaches have investigated how evoked activity propagates and is sustained through network interactions, PILI and ACFW quantify properties of the resulting perturbation trajectory, including its resilience and temporal memory. Extending the analysis window therefore allows longer-timescale changes in the post-stimulation response to be characterised, revealing aspects of response dynamics that may be missed when focusing solely on discrete early peaks.

The spectral and time-frequency findings further suggest that paired-pulse effects are not expressed uniformly across the evoked response. Differences between single-pulse and paired-pulse stimulation were evident in measures that reflect both cortical resilience and the temporal memory of oscillatory activity. This supports the view that LICI-related suppression may affect the organisation of the response across both frequency and time. Prior TMS-EEG studies have similarly shown that inhibitory effects can vary across evoked potentials and oscillatory features, rather than appearing as a uniform suppression effect (Rogasch et al., 2015).

### 4.3. Interpretation of PILI, ACFW, and LE

The three dynamical metrics captured various aspects of the perturbation response. PILI indexed the resilience of the post-stimulation response, providing a measure of how strongly the evoked response remained expressed over time. The measure was adapted from computational whole-brain modelling, in which PILI is utilized to evaluate how rapidly a system returns to its baseline state after an external perturbation and the time taken for network dynamics to return towards their baseline state (Jobst et al., 2021; Deco et al., 2018; Jobst et al., 2017). ACFW indexed temporal memory, reflecting how long the post-stimulation signal remained self-similar or autocorrelated. In complex systems theory, autocorrelation width is a foundational metric used to monitor critical slowing down and impending state transitions (Scheffer et al., 2009). These principles have been demonstrated in human neural recordings. Using long-term intracranial EEG, Maturana et al. (2020) showed that ACFW varied with seizure susceptibility and changed around transitions into seizure state, supporting the potential of autocorrelation-based measures to capture changes in temporal stability of human cortical activity (Maturana et al., 2020). TMS–EEG provides an experimental framework in which the cortical system can be directly perturbed and its subsequent response measured allowing dynamical properties such as temporal resilience and memory to be examined under controlled conditions. In the present dataset, PILI and ACFW were the measures most sensitive to differences between single-pulse and paired-pulse stimulation. This pattern suggests that the resilience and temporal memory of the post-stimulation response may be useful features for characterising LICI-related changes in cortical recovery.

The finding that ACFW distinguished stimulation conditions is particularly relevant because it demonstrates that inhibition alters the temporal features of the response beyond differences in response magnitude. Within a dynamical-systems framework, a shorter autocorrelation profile after paired-pulse stimulation indicates that temporal memory within the response decays more rapidly (i.e., more strongly damped effect). Together with the established pharmacological sensitivity of LICI to GABA_B_-mediated inhibition (Premoli et al., 2014), these findings suggest that controlled TMS–EEG perturbation may provide a non-invasive approach for examining how inhibitory conditioning modifies the temporal evolution of cortical activity.

Lyapunov exponent (LE) was included as an exploratory measure of dynamical stability. Lyapunov exponents have a long history in the analysis of neural dynamics. In animal models, particularly rodent models of temporal lobe epilepsy, the maximum Lyapunov exponent is a well validated tool for tracking the transition from chaotic interictal states to highly ordered ictal states. It provides a direct measure of how chaotic or predictable a neural system is (Mormann et al., 2007; Iasemidis et al., 1990). Theoretically, the Lyapunov exponent offers a distinct advantage by quantifying complex network interactions and fundamental dynamical stability that traditional linear metrics overlook (Iasemidis et al., 1990).

However, its weaker and less consistent performance in this dataset suggests that nonlinear stability metrics may be sensitive to methodological factors, including sample size, signal-to-noise ratio, embedding choices, preprocessing decisions, and the properties of scalp EEG. Animal studies typically use intracranial recordings with high signal to noise ratios, whereas the present study used scalp EEG, which is subject to volume conduction and lower spatial resolution (Stam, 2005). This does not mean that LE is not informative regarding the human brain, but rather that it must be interpreted cautiously. Future human studies using higher-density EEG, source-space reconstruction, or paradigms specifically designed to estimate nonlinear dynamics with larger samples may be better suited to evaluating whether LE provides reliable information about post-TMS stability.

Together, the metric pattern demonstrates that not all dynamical measures are equally informative for frontal LICI data. In the present dataset, PILI and ACFW showed greater sensitivity to paired-pulse modulation than LE, suggesting that measures capturing the resilience and temporal memory of the post-stimulation response may be well-suited for characterising perturbation-recovery dynamics using TMS-EEG, whereas LE may require further methodological validation. By adapting analytical tools traditionally reserved for animal models, intracranial electrophysiology and computational simulations to human perturbation data, this work offers a novel framework; moving beyond traditional amplitude measures towards characterising how the human cortical response evolves and recovers from targeted stimulation.

### 4.4. Open-source toolbox and clinical neurophysiology relevance

A practical contribution of this study is the implementation of these analyses in the open-source Brain Dynamic Metric Toolbox^1^. Standardised and transparent analysis tools are increasingly important for TMS-EEG, where preprocessing decisions, artefact handling, metric definitions, window choices, and statistical models can all influence results. Recent international guidance has emphasised that clinical translation of TMS-EEG will require reliable, reproducible, and interpretable outcome measures, supported by careful acquisition and analysis procedures (Tremblay et al., 2019; Ziemann et al., 2026).

The present toolbox contributes to this goal by providing a structured workflow for quantifying perturbation-recovery dynamics across temporal, spectral, and time-frequency domains. Although demonstrated here using frontal TMS-EEG data, the framework is not restricted to this paradigm. In principle, the same approach could be applied to other perturbation-response datasets, including electrical stimulation with EEG, intracranial stimulation recordings, or multimodal stimulation-recording studies. Its immediate value is therefore methodological: it supports reproducible calculation and comparison of dynamical response features that are otherwise difficult to quantify consistently across studies.

For clinical neurophysiology these measures may provide an additional dimension for characterising cortical dysfunction beyond conventional summaries of evoked-response magnitude. Alterations in excitation–inhibition balance and network regulation are implicated across a range of neurological and psychiatric disorders, raising the possibility that pathological differences may also be expressed in how cortical activity evolves and recovers following perturbation. The toolbox provides a reproducible framework for testing this hypothesis, although establishing the reliability, physiological specificity, and clinical sensitivity of individual dynamical metrics will require validation in larger independent and clinical cohorts.

### 4.5. Limitations and future directions

Several limitations should be considered. First, the sample consisted of healthy participants and was relatively small, so the results should be interpreted as a methodological demonstration rather than as validation of a clinical biomarker. The findings show that dynamical metrics can distinguish single-pulse and paired-pulse responses in this dataset, but further work is needed to establish reliability, normative ranges, clinical sensitivity, and diagnostic or prognostic utility.

Second, TMS-EEG is susceptible to both technical and physiological artefacts. Frontal stimulation is especially challenging because of muscle activity, auditory input, scalp sensation, and other sensory-evoked responses (Salo et al., 2020). Although masking noise and preprocessing reduce some sources of contamination, they do not eliminate sensory co-activation (Biabani et al., 2019). These sensory contributions are particularly important for the present analyses because they can extend beyond the immediate stimulation period and overlap with later TMS-evoked activity. Studies using sensory-control and sham conditions have demonstrated substantial sensory contributions from approximately 60 ms onwards, including within the prominent N100–P200 complex (Biabani et al., 2019; Conde et al., 2019; Freedberg et al., 2020; Gordon et al., 2023). Within the present 950 ms analysis framework, these effects therefore occur relatively early in the post-stimulation trajectory and may influence subsequent measures of response resilience and temporal memory.

This issue requires particular consideration when comparing single-pulse and paired-pulse stimulation. Paired-pulse trials contain an additional conditioning stimulus and therefore differ from single-pulse trials not only in cortical inhibitory engagement, but also in the number and temporal sequence of associated auditory and somatosensory events. Sensory-evoked activity associated with the conditioning pulse may consequently overlap with the response to the subsequent test pulse. As PILI and ACFW characterise the resilience and temporal memory of the post-stimulation signal, sensory activity occurring within the extended response window could contribute to these measures and cannot be separated from cortical perturbation dynamics on the basis of the present data alone. The dynamical framework therefore does not resolve sensory confounding; rather, it provides a structured way of quantifying how the measured post-stimulation response evolves over time. Future studies incorporating sensory-matched single-and paired-pulse sham conditions will be important for determining the extent to which changes in PILI and ACFW reflect cortical inhibitory processes rather than differences in sensory co-activation.

Third, temporal inference depends on the selected windowing scheme. The use of fixed 150 ms windows provides a transparent and interpretable analysis structure but imposes discrete boundaries on a continuous response. Future analyses should assess whether the observed effects are robust across alternative bin widths, overlapping sliding windows, or adaptive temporal approaches. Such analyses would help determine whether the findings reflect stable recovery features or are partly dependent on window placement.

Finally, the biological interpretation of the metrics requires further validation. Reduced PILI and ACFW after paired-pulse stimulation are consistent with inhibitory modulation of the evoked response, but they do not directly identify the underlying synaptic or neurochemical mechanisms. Future studies combining TMS-EEG dynamical metrics with pharmacological manipulation, magnetic resonance spectroscopy, source modelling, sham-controlled sensory conditions, and clinical outcome measures will be important for clarifying their physiological and translational significance.

## 5. Conclusion

This study demonstrates that single-pulse and paired-pulse frontal TMS-EEG responses can be differentiated using dynamical metrics that capture complementary features of cortical perturbation recovery. PILI and ACFW were particularly sensitive to LICI-related changes in cortical resilience and temporal memory, whereas LE provided more exploratory information about dynamical stability. These findings support a broader view of paired-pulse inhibition as altering the temporal dynamics of the evoked response, rather than expressed solely as a reduction in amplitude at selected latencies or electrodes.

By implementing these analyses in an open-source toolbox, the present work provides a reproducible framework for quantifying perturbation-recovery dynamics in TMS-EEG. The approach may be useful for re-analysing existing datasets and for designing future studies of cortical inhibition, recovery, and treatment response. Further validation in larger, well-controlled cohorts will be needed before these metrics can be interpreted as clinical biomarkers, but the present findings provide a foundation for studying how inhibitory perturbations shape cortical recovery dynamics in health and disease.

## 6. Acknowledgements

We sincerely thank Dianne Sheppard and the late John Bradshaw for their guidance and contributions to the design of the original study, from which the data for the present analysis were derived. We also thank Nigel Rogasch, Jerome Maller, Neil Bailey, and Karyn Richardson for their support and contributions to participant recruitment and data acquisition during the original study. Finally, we thank Christian Davey from the Statistical Consulting Platform at the University of Melbourne for his valuable advice and assistance with the statistical analyses.

## 7. Funder Information Declared

- National Health and Medical Research Council, APP2033596
- Australian Research Council, DP240102899

## Footnotes

1 https://github.com/qinwayne/BrainDynamicMetricToolbox

